# Recent secondary contacts, background selection and variable recombination rates shape genomic diversity in the model species *Anolis carolinensis*

**DOI:** 10.1101/352922

**Authors:** Yann Bourgeois, Robert P. Ruggiero, Joseph D. Manthey, Stéphane Boissinot

## Abstract

Gaining a better understanding on how selection and neutral processes affect genomic diversity is essential to gain better insights into the mechanisms driving adaptation and speciation. However, the evolutionary processes affecting variation at a genomic scale have not been investigated in most vertebrate lineages. Previous studies have been limited to a small number of model species, mostly mammals, and no studies have investigated genomic variation in non avian reptiles. Here we present the first population genomics survey using whole genome re sequencing in the green anole (*Anolis carolinensis*). This species has emerged as a model for the study of genomic evolution in squamates. We quantified how demography, recombination and selection have led to the current genetic diversity of the green anole by using whole-genome resequencing of five genetic clusters covering the entire species range. The differentiation of green anole’s populations is consistent with a northward expansion from South Florida followed by genetic isolation and subsequent gene flow among adjacent genetic clusters. Dispersal out-of-Florida was accompanied by a drastic population bottleneck followed by a rapid population expansion. This event was accompanied by male-biased dispersal and/or selective sweeps on the X chromosome. We show that the combined effect of background selection and recombination rates is the main contributor to the genomic landscape of differentiation in the anole genome. We further demonstrate that recombination rates are positively correlated with GC content at third codon position (GC3) and confirm the importance of biased gene conversion in shaping genome wide patterns of diversity in reptiles.

## Introduction

Nucleotide variation along a DNA sequence results from the interactions between multiple processes that either generate new alleles (e.g. recombination, mutation) or affect the fate of these alleles in populations (e.g. selection, demography and speciation). The variable outcome of these interactions along the genome can result in heterogeneous patterns of diversity and divergence at both intra- and inter-specific scales (Cruickshank and Hahn 2014; Roux et al. 2014; Seehausen et al. 2014; Wolf and Ellegren 2016). Given their importance, quantifying these processes has been at the core of evolutionary genomics for the last decade. With the advent of next-generation sequencing and the continuous development of novel analytical tools, it has become possible to properly quantify the impact of recombination (Booker et al. 2017; Kawakami et al. 2017), selection (Barrett et al. 2008; Mullen and Hoekstra 2008) and demographic history (Gutenkunst et al. 2009; Excoffier et al. 2013; Roux et al. 2016) on diversity patterns in several vertebrates.

Ultimately, such investigations have the power to answer outstanding biological questions such as the role of sex chromosomes, the nature of reproductive barriers or the timing of gene flow during the process of speciation (Wolf and Ellegren 2016). However, genomic patterns of variation retrieved from genome scans need to be interpreted with caution. For example, recent years have seen a growing interest for the so-called “genomic islands of divergence”, those genomic regions that harbor high differentiation between species or populations (Ravinet et al. 2017). This pattern was at first interpreted as evidence for genomic islands resisting gene flow and introgression but was often based on the examination of relative differentiation statistics, such as F_ST_ (Cruickshank and Hahn 2014). However, further investigation using absolute measures of divergence demonstrated an important role of selection reducing diversity in regions of low recombination. This questioned the emphasis put on genomic barriers to gene flow in heterogeneous divergence along the genomes of incipient species (Cruickshank and Hahn 2014). The difficulties in interpreting genome-wide patterns of diversity and differentiation can nonetheless be alleviated by combining information from different methods to properly take into account the factors that may produce similar distributions for the statistics of interest (Cruickshank and Hahn 2014; Ravinet et al. 2017). Recent advances in model-fitting of demographic scenarios incorporating heterogeneity in selection and gene flow along genomes have contributed to a better assessment of these evolutionary forces on genomic diversity of wild populations (Roux et al. 2014; Christe et al. 2016; Roux et al. 2016).

Despite the absolute necessity to increase sampling across the tree of life (Abzhanov et al. 2008), it remains that some clades are still poorly studied in a population genomics context. It is thus unclear if patterns observed in the most commonly studied organisms (e.g. human and mouse) apply widely. Thorough analyses of the factors affecting genome diversity at the peri-specific level have been performed in a small number of vertebrate species (see Ellegren *et al.*, 2012; Sousa *et al.*, 2013; Poelstra *et al.*, 2014; Booker *et al.*, 2017; Han *et al.*, 2017; Kawakami *et al.*, 2017) but several major clades of vertebrates have not been investigated at all. This need for higher resolution in genomic studies has been recently highlighted with several comments urging to “scale-up” the sequencing effort in studies of evolutionary radiations around the species level (Ellegren 2014; de la Harpe et al. 2017). Among those are the non-avian reptiles, a speciose group of vertebrates that harbor a wide diversity of morphology and adaptation. To fill this gap in our knowledge, we decided to perform a study on genome-wide variation in the green anole *(Anolis carolinensis),* a model species for behavior, physiology and comparative genomics (Tollis et al. 2012; Wade 2012; Tollis and Boissinot 2014; Manthey et al. 2016).

The green anole is the first non-avian reptile for which we have a complete genome sequence (Alföldi et al. 2011) and its genetic structure is relatively well known (Tollis et al. 2012; Tollis and Boissinot 2014; Campbell-Staton et al. 2016; Manthey et al. 2016; Ruggiero et al. 2017). Since a reference genome is available, whole-genome resequencing of anoles populations would be an opportunity to better understand the drivers and constraints that act on species radiations at a resolution that was not allowed by the genetic datasets used in previous studies.

The green anole colonized Florida from Cuba (Glor et al. 2005; Campbell-Staton et al. 2012; Tollis et al. 2012; Tollis and Boissinot 2014; Manthey et al. 2016) between 6 and 12 million years ago and populations in Florida likely diverged in allopatry on island refugia before secondary contact due to sea-level oscillations during the Pleistocene. Colonization of the rest of North America seems to be more recent, with two clades having probably expanded in the last 500,000 years (Manthey et al. 2016). This recent radiation makes the green anole a suitable model to study how purifying selection, recombination, and barriers to gene flow shape genomic diversity in a reptile.

In addition, despite the availability of a genomic reference, our knowledge of the fundamental processes that drive genome evolution over long timescales remains limited in squamates. For example, it has been suggested that the genome of the green anole lacks GC isochores, an unusual feature in vertebrates (Alföldi et al. 2011; Fujita et al. 2011). These initial studies have suggested that uniform recombination rates or lack of biased gene conversion increasing GC content in regions of high recombination (Marais 2003) might explain this seemingly homogeneous landscape. However, the claim of homogeneity was recently rebutted (Costantini et al. 2016), though the GC content of green anoles does seem to be more uniform than in other vertebrates (Alföldi et al. 2011; Figuet et al. 2014; Costantini et al. 2016). Nevertheless, high heterogeneity of GC content at the third codon position (GC3) in the green anole genome strongly suggests biased gene conversion and heterogeneous recombination rates (Figuet et al. 2014; Costantini et al. 2016). Clarifying such controversies would benefit from a genome-wide analysis of GC content and recombination.

Here we present results obtained from whole-genome resequencing of five genetic clusters of the green anole. We provide for the first time a detailed assessment of the multiple factors that are likely to impact the green anole’s genetic diversity at a genome-wide scale. We demonstrate that the combined effects of regional variation in recombination rate, background selection, and migration are responsible for the heterogeneous genomic landscape of diversity and divergence in the green anole.

## Results

### Statistics for whole genome resequencing

Twenty-seven green anoles (*Anolis carolinensis*) sampled across the species’ range and covering the five genetic clusters identified in previous analyses (Tollis and Boissinot 2014; Manthey et al. 2016) were chosen for whole-genome resequencing. We also included two samples from the closely-related species *A. porcatus* and *A. allisoni* as outgroups. Sequencing depth was comprised between 7.22X and 16.74X, with an average depth of 11.45X (Table S1). 74,920,333 variants with less than 40% missing data were retained after the first round of filtering (Methods).

### Population structure and nucleotide variation reveal a decrease of diversity in northern populations

Using more than 6,500 unlinked SNPs with less than 20% missing data of all green anole samples, DAPC identified k=5 as the most likely number of genetic clusters (Figure 1A). These groups were consistent with the clusters identified in previous genetic studies (Tollis et al. 2012; Tollis and Boissinot 2014; Manthey et al. 2016). Possible introgression from Carolinas was observed for two Gulf Atlantic individuals (Figure 1A). A maximum-likelihood phylogeny estimated in RAxML and a network analysis of relatedness in Splitstree further supported this clustering (Figure 1B, C). Results closely matched previous findings, with South Florida (SF) being the sister clade of all other groups. The two northernmost clusters, Gulf Atlantic (GA) and Carolinas (CA), clustered together in the RAxML phylogeny. Eastern Florida (EF) constituted a paraphyletic group in the phylogeny in which GA and CA were nested. This is likely due to incomplete lineage sorting induced by the high and constant effective population sizes of populations from Florida (see below), or to ongoing or recent gene flow resulting in the inclusion of loci with different coalescence times. At last, the Western Florida (WF) cluster was basal to all other groups except South Florida.

**Figure 1.**
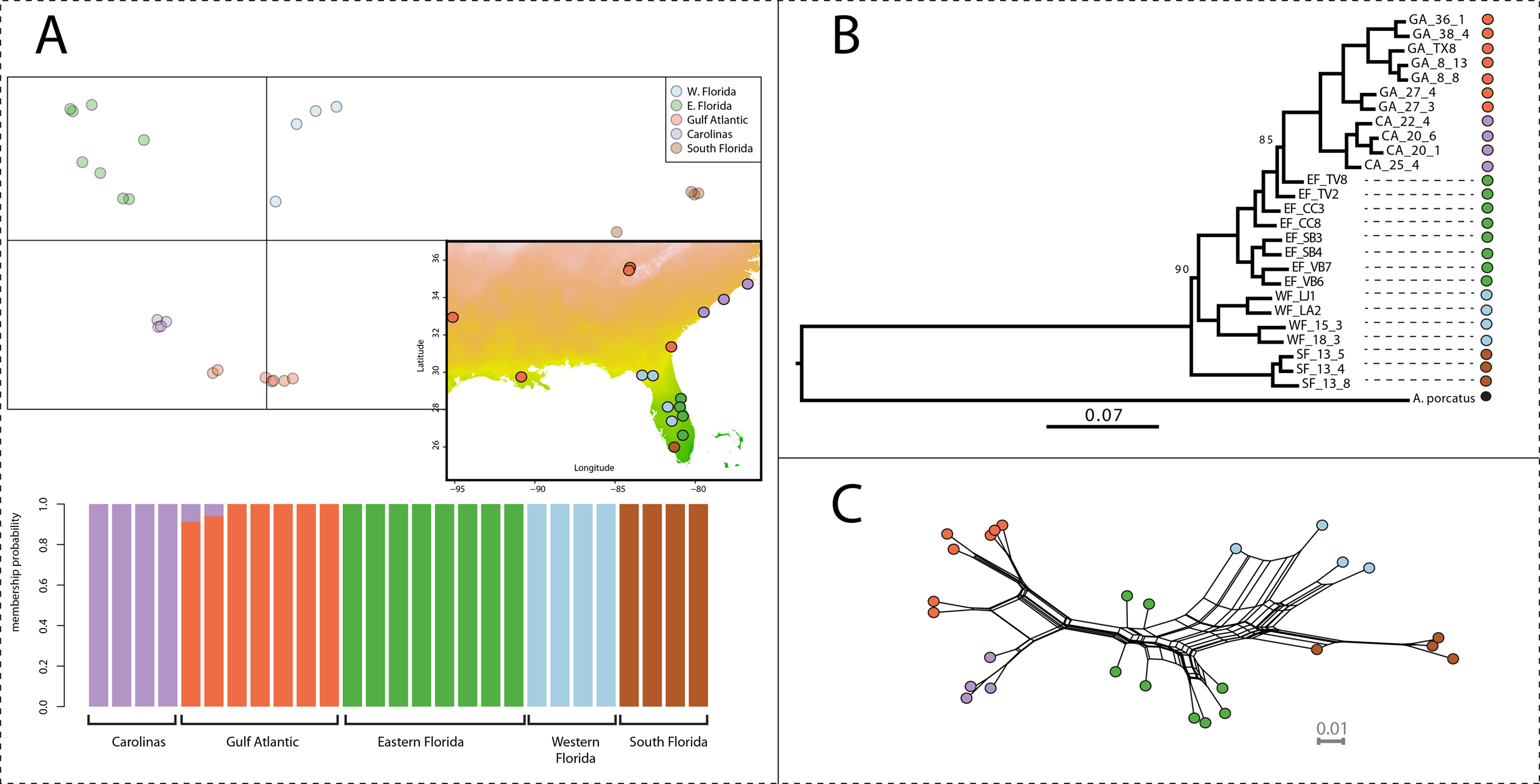
Genetic structure in *Anolis carolinensis* from whole-genome SNP data. A: Results from the DAPC analysis highlighting the five clusters inferred from the analysis of ~6,500 SNPs thinned every 10kb and with less than 20% missing data. The map reports the coordinates of the localities used in this study and the genetic clusters they belong to. B: RAxML phylogeny based on one million SNPs randomly sampled across the genome. All 100 bootstrap replicates supported the reported topology, except for two nodes with support of 90 and 85. One individual from South Florida was removed due to a high rate of missing data. C: Network representation of the relatedness between samples as inferred by Splitstree v4. Color codes match those in parts A and B.

Nucleotide diversity was the lowest in GA and CA (Table 1) despite the large geographic area covered by these two genetic clusters. Tajima’s D values ranged between -0.8 (WF) and 0.14 (GA). Positive Tajima’s D values suggest recent population contraction, while negative Tajima’s D are expected in the case of recent population expansion (Tajima 1989). No evidence for strong recent bottlenecks could be observed, although northern clusters (CA and GA) displayed the highest average Tajima’s D.

**Table 1.**
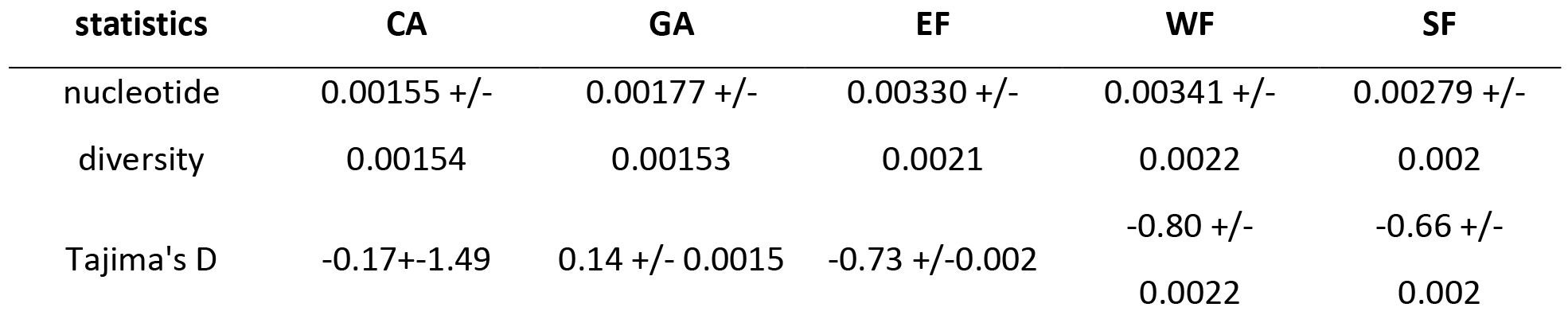
Diversity and Tajima’s D (+/- s.d.) for each of the five genetic clusters, averaged over non-overlapping 5kb windows across the genome.

| statistics | CA | GA | EF | WF | SF |
| --- | --- | --- | --- | --- | --- |
| nucleotide | 0.00155 +/- | 0.00177 +/- | 0.00330 +/- | 0.00341 +/- | 0.00279 +/- |
| diversity | 0.00154 | 0.00153 | 0.0021 | 0.0022 | 0.002 |
| Tajima's D | -0.17+/-1.49 | 0.14 +/- 0.0015 | -0.73 +/--0.002 | -0.80 +/-<br>0.0022 | -0.66 +/-<br>0.002 |

### Recent population expansion and male-biased sex-ratios in Northern populations

We used the whole set of filtered SNPs with less than 40% data to infer past changes in effective population sizes (Ne) without any *a priori* demographic model with SMC++ (Figure 2A). All populations from Florida showed rather stable demographic trajectories, with some evidence for population expansion in EF and WF. Assuming a mutation rate of 2.1×10^−10^/bp/year (Tollis and Boissinot 2014), population sizes were in the range of 500,000 to 5,000,000 individuals for each population, in accordance with previous analyses based on target capture markers (Manthey et al. 2016). Northern populations (CA and GA) showed a clear signature of expansion starting between 200,000 and 100,000 years ago, following a bottleneck that started between 500,000 and 1,000,000 years in the past. We also estimated the splitting times between the different groups but since this model assumes no gene flow after the split, the estimates are likely to be biased toward the present. The split between GA and CA occurred shortly before these populations expanded, in accordance with the previously proposed hypothesis of double colonization following the Gulf and Atlantic coasts (Tollis and Boissinot 2014). In Florida, divergence events took place between 3 and 2 million years ago. The relative order of splitting events was consistent with the topology obtained from our phylogeny and previous studies.

**Figure 2.**
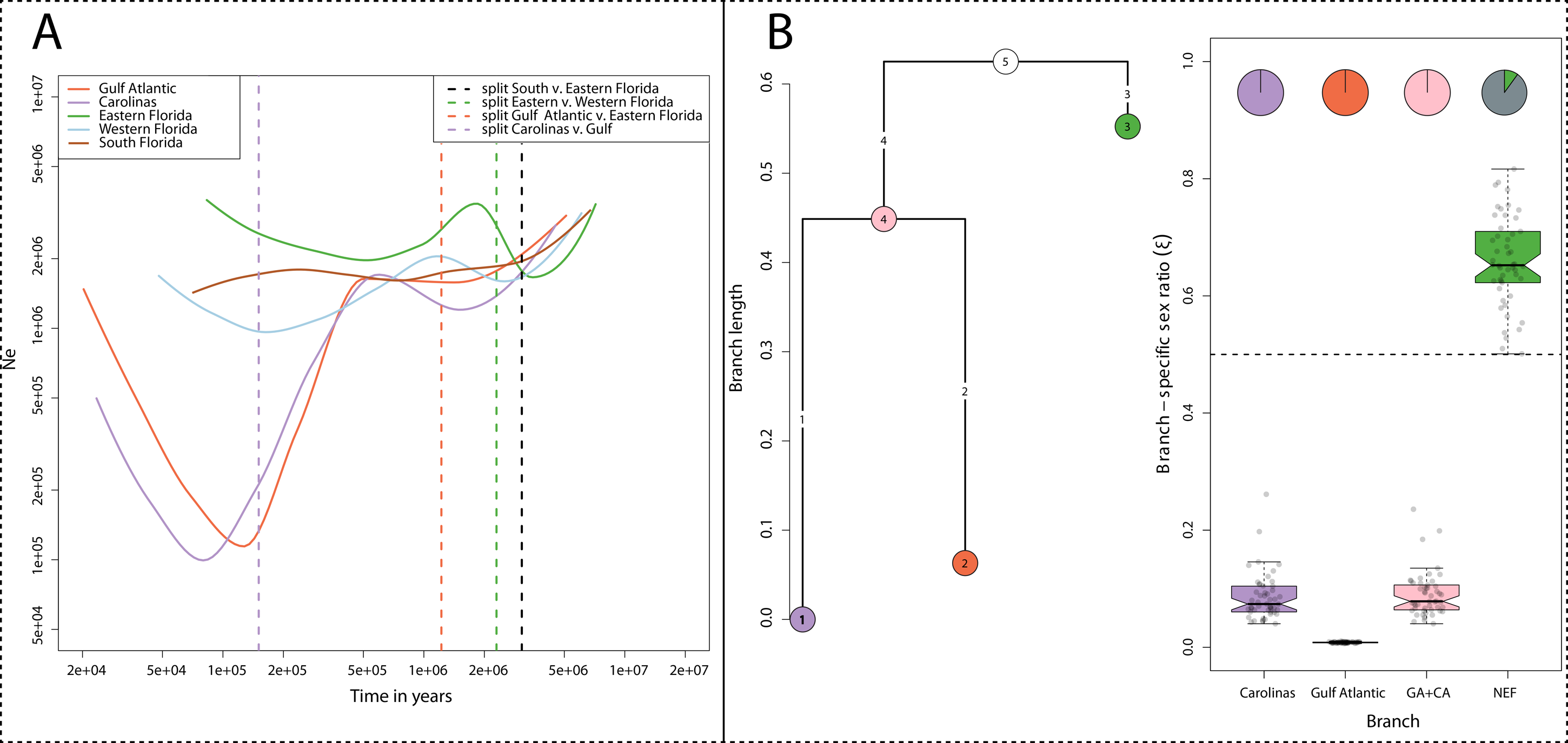
Variation in effective population sizes with time and comparison of drift between autosomes and sex-linked scaffolds. A: Reconstruction of past variations in effective population sizes (Ne) inferred by SMC++. Dashed vertical lines correspond to the estimated splitting times between the five genetic clusters previously inferred. We assume a mutation rate of 2.1×10^−10^/bp/generation and a generation time of one year. B: Average branch lengths obtained from autosomal data and effective sex-ratios (ξ) inferred from KIMTREE. A set of 5,000 autosomal and 5,000 sex-linked markers were randomly sampled to create 50 pseudo-replicated datasets on which the analysis was run. The analysis was run on the three most closely related populations. Pie charts indicate the proportion of replicates for which we observed significant support (S_i_<0.01) in favor of a biased sex-ratio.

Anoles are an important model for studying behavior and conflicts between and within sexes (Johansson et al. 2008). We tested whether the recent colonization of new and possibly suboptimal habitats could lead to a shift in the reproductive dynamic of anoles or sexual selection (Figure 2B). We built a population tree and quantified genetic signatures of biased sex-ratio with the algorithm implemented in KIMTREE. We focused on the three populations that diverged most recently, GA, CA and EF. Note that the length of branch i (τ_i_) represents time in generations (t_i_) scaled by the effective population size for this branch such as τ_i_ = t_i_/2N_e,i_ (Clemente et al. 2018). Branch lengths were particularly high for the CA and GA lineages compared to EF, as expected in the case of stronger drift (Figure 2B). This is in line with their smaller effective population sizes and the bottleneck inferred by SMC++. We found evidence for a strongly male-biased effective sex-ratio (ESR) in CA and GA, but not EF which was slightly female-biased. Indeed, nucleotide diversity was substantially more reduced at sex-linked scaffolds in GA than in EF when compared to autosomal diversity (Sup. Fig. 2). Note that sex-ratios are the proportion of females effectively contributing to the gene pool along each branch of the tree and should not be interpreted directly in terms of census size. The GA cluster displayed the strongest bias, with an estimated ratio of less than one female for 100 males, suggesting strong sex-bias in the founding population or strong male-biased dispersal during population expansion. The CA cluster and the inner branch leading to CA and GA showed a ratio of approximately ten females for 100 males. All 50 replicates displayed a high support for a male-biased sex-ratio in CA and GA, while only 5 replicates supported a female-biased sex-ratio in EF (i.e. the Markov chain almost systematically explored sex-ratios above 0.5 in only 5 replicates).

### Secondary contact and gene flow have homogenized green anole populations

We tested whether secondary contact may have played a role in shaping the genomic landscape of differentiation in green anoles (Figure 3). We compared a set of 34 divergence scenarios, allowing gene flow and effective population sizes to vary with time and across loci. Briefly, heterogeneity in gene flow (suffix 2M2P) was implemented by dividing the site frequency spectrum into three sets of loci with proportions 1-P_1_-P_2_, P_1_ and P_2_. The first set (1-P_1_-P_2_) was modelled with all parameters from the base model. The two other sets were modelled with no gene flow towards population 1 (P_1_) or population 2 (P_2_) and represent genomic islands resisting gene flow in populations 1 and 2 respectively. To simulate the reduction in diversity expected under purifying selection at linked, non-recombinant (nr) sites, two sets of loci were modelled at frequencies 1-nr and nr (suffix 2N). The first set was modelled with all parameters from the base model, the other with the same parameters but with effective population sizes reduced by a background selection factor (bf).

**Figure 3.**
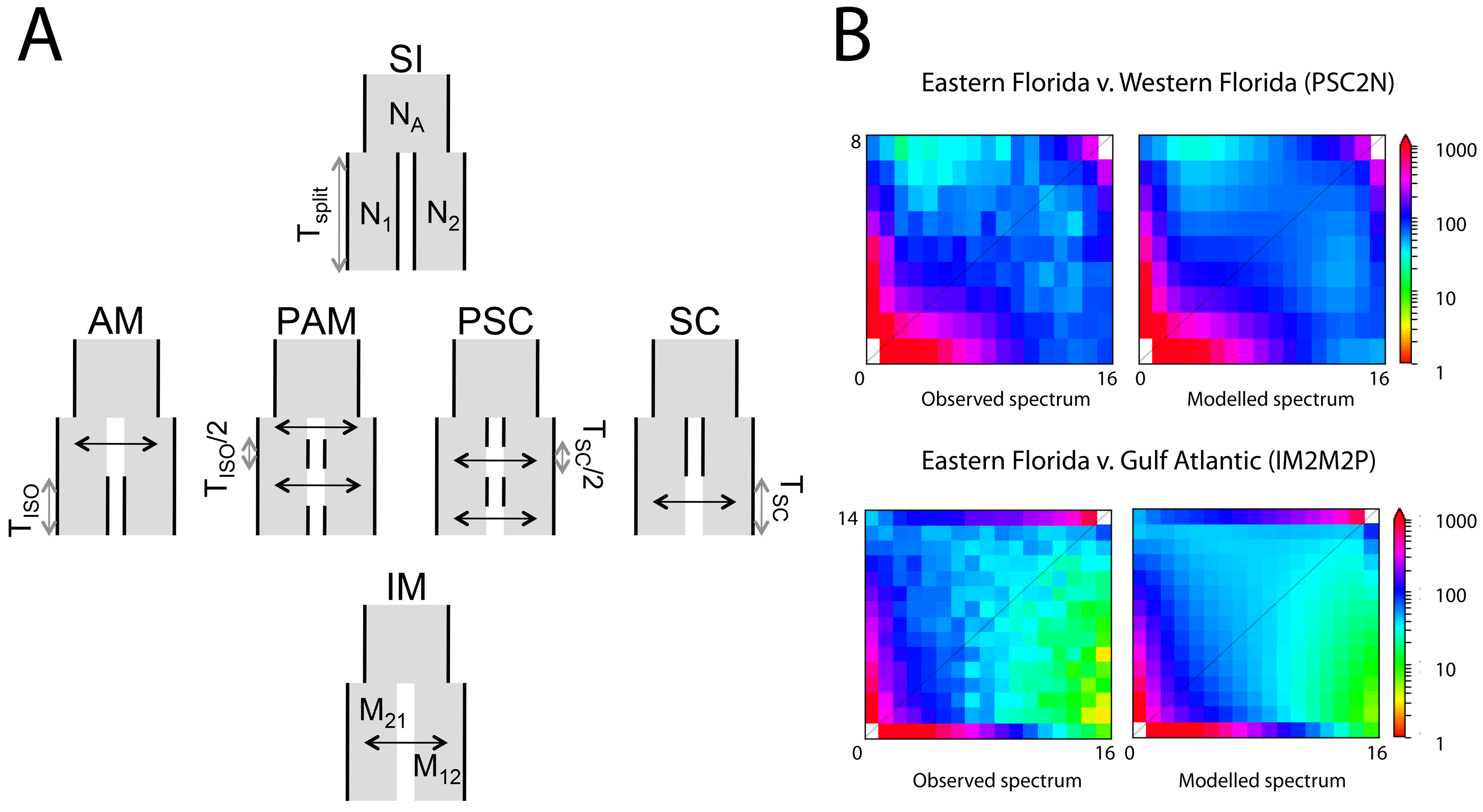
A: Graphic description of the 6 categories of ∂a∂i models tested over pairs of green anole genetic clusters. Each model describes a scenario where two populations diverge from an ancestral one, with varying timing and strength of gene flow after their split. SI: Strict Isolation; AM: Ancestral Migration where populations first exchange gene flow then stops T_iso_ generations ago; PAM: Ancestral migration with two periods of contact lasting T_iso_/2 generations; SC: Secondary Contact where populations still exchange gene flow at present time; PSC: Secondary contact with two periods of contact lasting T_sc_/2 generations; IM: Isolation with constant migration and no interruption of gene flow. Reproduced with the authorization of Christelle Fraïsse. B: Fitting of the best models for the EF (N=16) v. GA (N=14) and EF v. WF (N=8) comparisons. Both models fit the observed datasets as indicated by the similar spectra between observation and simulation. The “2N” suffix means that background selection was added to the base model by modelling heterogeneous effective population sizes across loci. The “2M2P” suffix means that heterogeneity in gene flow was incorporated into the model. The “ex” suffix means that exponential population size change was introduced in the base model.

Strict-isolation models (SI) consistently displayed the lowest likelihood, clearly supporting a role for gene flow in homogenizing green anoles genomes. For the comparison between EF and WF, models including heterogeneous population sizes performed better than models with heterogeneous gene flow. Among scenarios with gene flow, secondary contact with one and two periods of gene flow (SC and PSC) often received the highest support (Figure 3B, Figure 4). Parameters estimated from the best models are shown in Table 2. There was no substantial gain in likelihood when adding expansion to scenario of two secondary contacts with background selection (PSC2N), and models with heterogeneous migration displayed lower likelihood. The PSC2N model supported a scenario where about nr = 65% of the genome was affected by background selection, suggesting a rather large effect of low recombination and purifying selection on diversity. These Eastern and Western Floridian genetic clusters experienced long periods of isolation lasting about 2 million years, followed by periods of secondary contact lasting approximately 125,000 years in total.

**Figure 4.**
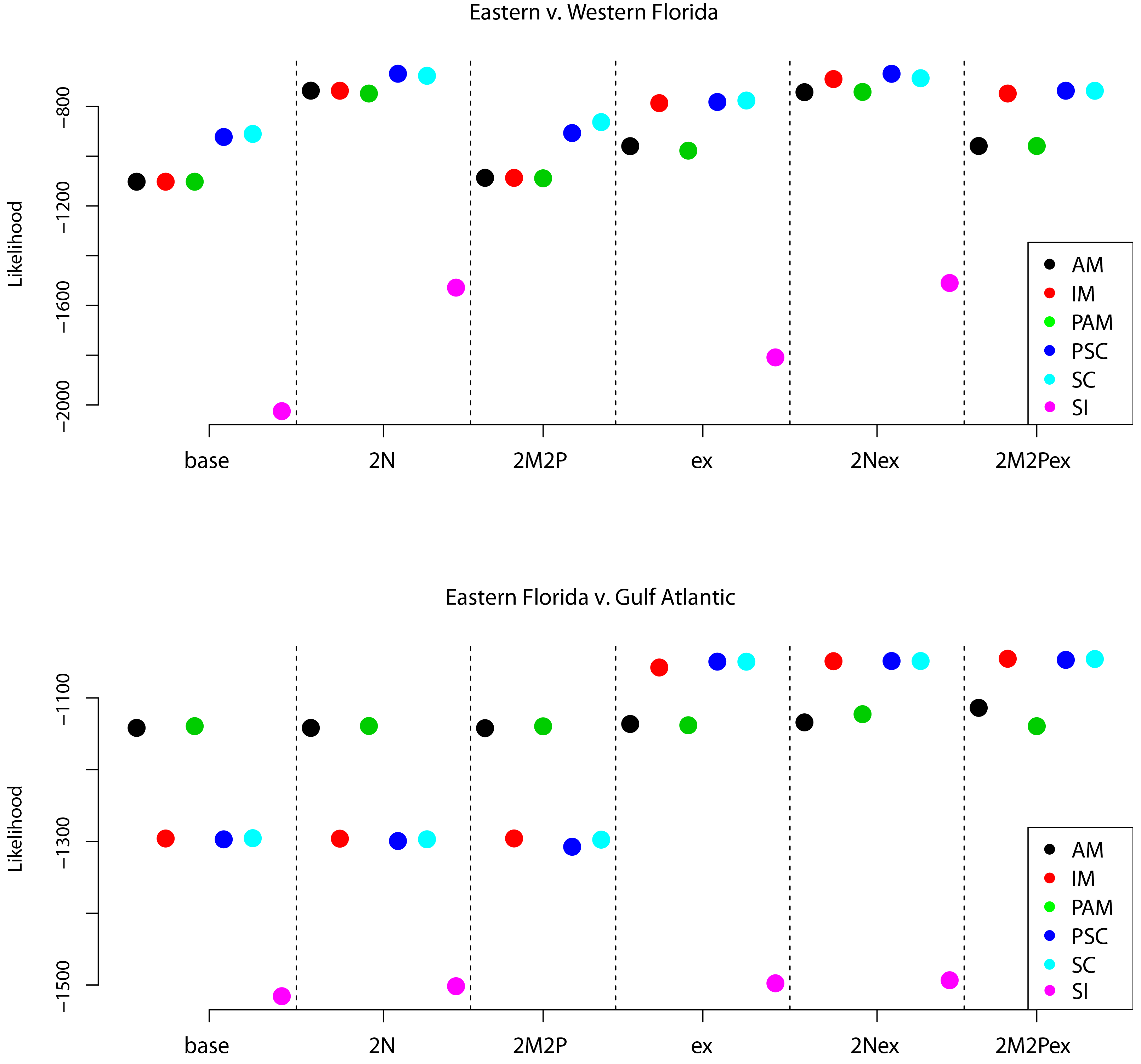
Likelihoods obtained for the 34 ∂a∂i models in the EF v. GA and EF v. WF comparisons. Higher likelihoods suggest better support for a given model. Complexity was added to the models described in Figure 3A by including various combinations of population expansion (prefix ‘ex’), heterogeneous asymmetric migration rates (suffix ‘2M2P’) and heterogeneous effective population size (suffix ‘2N’) among loci.

**Table 2.** Summary of best-supported demographic models. PSC2N: Secondary contact with two periods in isolation and heterogeneous effective population sizes across the genome. SCex: Secondary contact with an episode of population expansion following secondary contact. SC2M2P and IM2M2P: Models of secondary contact and constant gene flow with heterogeneous migration rates along the genome. nr: proportion of the genome displaying an effective population size of bf times the population size displayed by the remaining 1-nr fraction not affected by background selection. O: proportion of sites for which the ancestral state was correctly inferred. P1 and P2 are the proportion of sites resisting gene flow in populations 1 and 2. Tiso: total time spent in isolation. For the PSC model, populations are isolated twice in their history for Tiso/2 generations and are connected twice for Tsc/2 generations (see Figure 3A). Tsc: time during which stable populations stay connected. Tscg: Time since population size change (with gene flow). The total time during which populations were connected is Tscg+Tsc. For each model, the first line shows the set of best estimates, and the second the standard deviation obtained from 100 bootstrap replicates.

| Comparison<br>(Pop. 1 v Pop. 2) |  | Model | theta | Ne1 | Ne2 | m2->1 | m1->2 | Tiso | Tsc | Tscg | nr | bf | P1 | P2 | O | Likelihood | AIC |
| --- | --- | --- | --- | --- | --- | --- | --- | --- | --- | --- | --- | --- | --- | --- | --- | --- | --- |
| WF v EF |  | PSC2N | 2091300 | 5482659 | 5731754 | 4.60E-06 | 1.64E-06 | 2135030 | 126496 | NA | 0.61 | 0.24 | NA | NA | 0.98 | -668.32 | 1354.67 |
| +/- s.d. |  | PSC2N |  | 761890 | 804877 | 3.40E-06 | 1.18E-06 | 64101 | 104327 | NA | 6.10E-02 | 2.04E-02 | NA | NA | 1.59E-03 |  |  |
| GA v EF |  | SCex | 2156641 | 382017 | 7731535 | 3.60E-07 | 4.85E-07 | 571481 | 125 | 1369541 | NA | NA | NA | NA | 0.97 | -1049.45 | 2118.90 |
| +/- s.d. |  | SCex |  | 8816 | 1068901 | 9.42E-08 | 4.63E-08 | 411287 | 1078789 | 114165 | NA | NA | NA | NA | 2.54E-03 |  |  |
| GA v EF |  | SC2M2Pex | 2156641 | 387945 | 8066417 | 4.42E-07 | 5.40E-07 | 1 | 664554 | 1399941 | NA | NA | 0.83 | 0.93 | 0.97 | -1045.70 | 2115.40 |
| +/- s.d. |  | SC2M2Pex |  | 10044 | 951413 | 3.81E-08 | 1.62E-08 | 2003745 | 1195746 | 63302 | NA | NA | 0.10 | 0.02 | 2.41E-03 |  |  |
| GA v EF |  | IM2M2Pex | 2156641 | 375534 | 7936973 | 5.54E-07 | 5.30E-07 | NA | 647269 | 1380877 | NA | NA | 0.73 | 0.93 | 0.97 | -1045.37 | 2112.74 |
| +/- s.d. |  | IM2M2Pex |  | 9146 | 336313 | 1.25E-08 | 3.13E-08 | NA | 152996 | 45278 | NA | NA | 0.04 | 0.03 | 1.59E-03 |  |  |

For the comparison between GA and EF, we confirmed a smaller effective population size in GA compared to EF (about 20 times smaller). The model with the smallest AIC was the IM2M2Pex model, followed by models of secondary contact (PSCex, SCex, and SC2M2Pex). We therefore present results obtained for several representative models (Table 2). All models supported a scenario with extensive gene flow, with high uncertainties for the time spent in isolation for secondary contact models (SC). Models with the highest likelihood and AIC incorporated genomic barriers to gene flow in GA, with approximately 20-30% of loci resisting introgression from Florida and less than 10% resisting gene flow from GA.

### Recombination and purifying selection shape genome composition and allele frequencies

Secondary contacts are often associated with the emergence of genomic islands resisting gene flow, that display higher differentiation than regions that have been homogenized. The diversity of such islands is also higher, as they diverged and accumulated mutations before gene flow resumed. On the other hand, purifying selection at linked sites can also generate genomic islands, as it reduces diversity and lead to an increase of relative measures of differentiation (Cruickshank and Hahn 2014). Some of the best supported models in ∂a∂i suggested a widespread impact of background selection in Florida, reducing diversity at linked sites over ~60% of the genome. We therefore tested the role of low recombination in shaping the genomic landscape of diversity and differentiation in green anoles in a context of secondary contact. Recombination rates estimated by LDHat in the EF cluster were highly heterogeneous along chromosomes, with stronger recombination rates at the tips and towards centromeres, though they dropped at the immediate vicinity of the latter (Figure 5). This pattern was supported by the Rozas’s ZZ statistic, suggesting stronger linkage disequilibrium in the middle of chromosomes arms.

**Figure 5.**
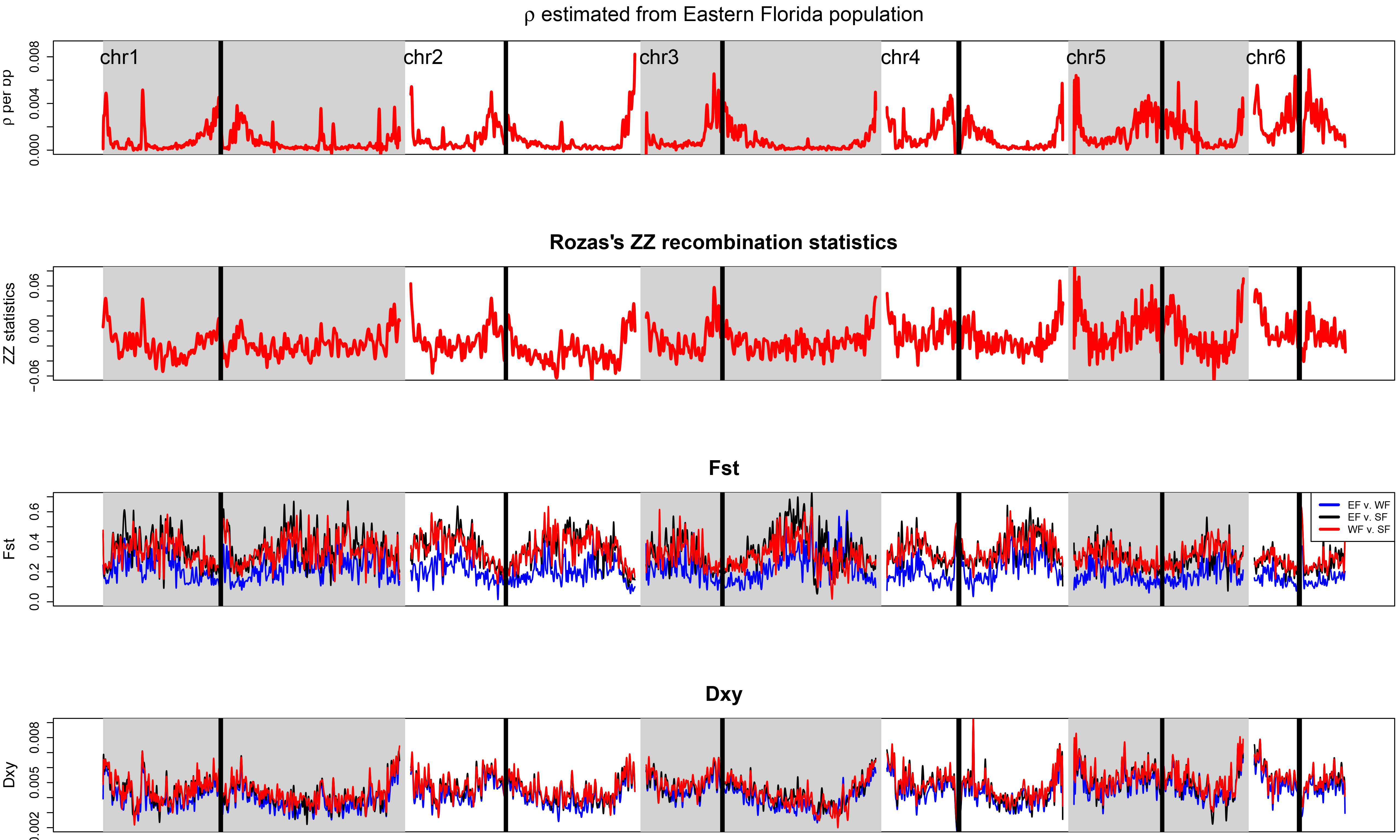
Summary statistics for recombination and differentiation along chromosomes. ρ= 4*Ne*r, with r the recombination rate per bp and per generation and Ne the effective population size for the EF cluster. Rozas’s ZZ is a measure of linkage disequilibrium positively correlated to intragenic recombination. FST and dXY are relative and absolute measures of differentiation that are correlated with the amount of shared heterozygosity and coalescence time across populations respectively. We present differentiation for the three genetic clusters having diverged for the longest time period. Statistics were averaged over non-overlapping 5kb windows and a smoothing line was fit to facilitate visual comparison. Repetitive centromeric regions that are masked from the green anole genome are highlighted by black rectangles.

We observed higher relative differentiation (measured by F_ST_) in regions of low recombination (Spearman’s rank correlation test, all P-values < 2.2×10^−16^; Figure 5, Figure 6). The correlation was however opposite for measures of absolute differentiation (d_XY_), a statistics directly related to diversity and average age of alleles across populations (Cruickshank and Hahn 2014). These correlations are consistent with selection reducing heterozygosity in regions of low recombination, and further support the ∂a∂i models of heterogeneous effective population sizes along the genome.

**Figure 6.**
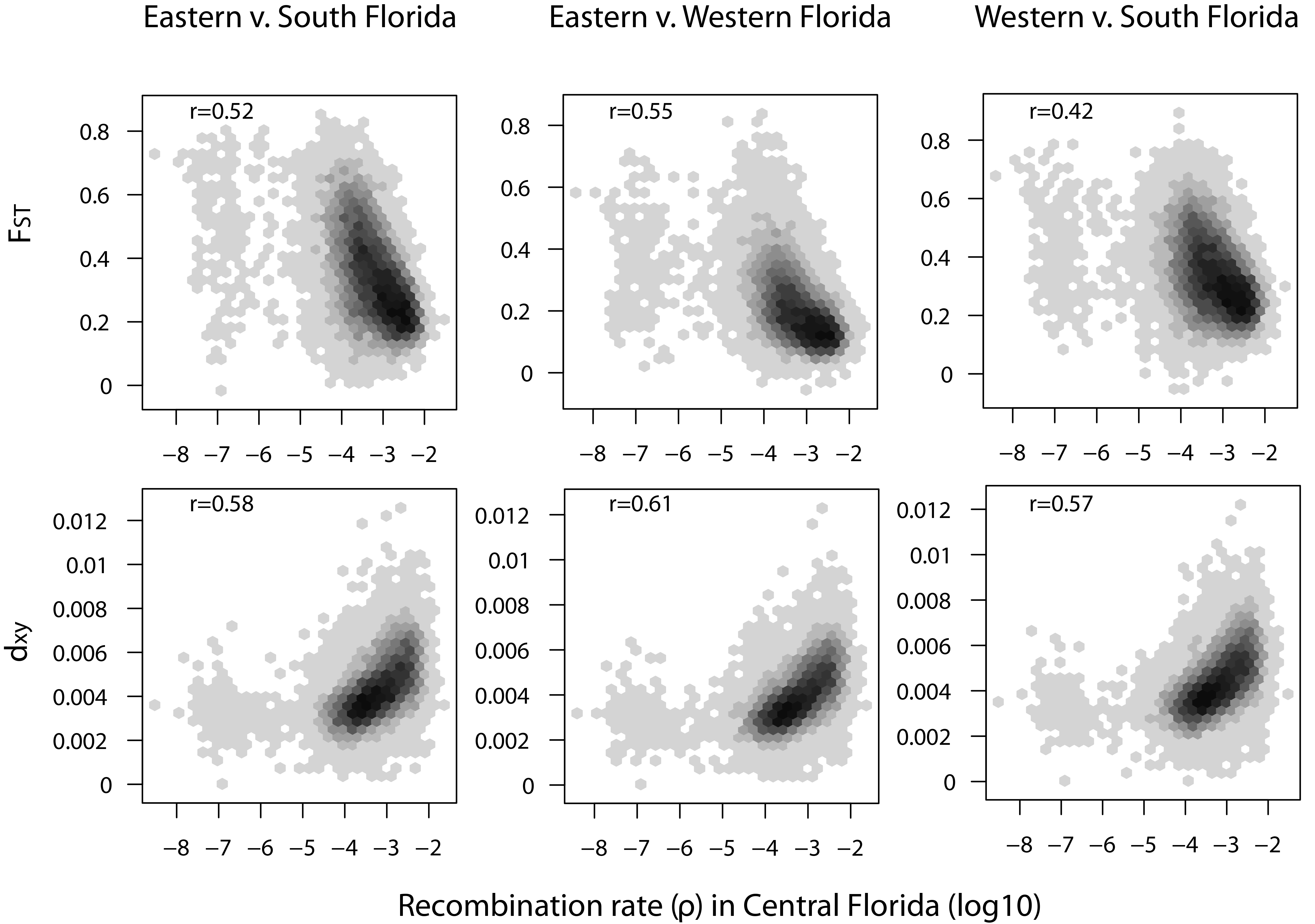
Correlations between ρ, FST, and dXY at the genome scale. Statistics were averaged over non-overlapping 5kb windows. Spearman’s ρ coefficients (r) are indicated on the graphs.

We assessed whether biased gene conversion had an impact on nucleotide composition in the green anole by testing for correlation between recombination and GC content in coding DNA sequences (CDS). We did observe a significant correlation between GC content and recombination rates at all three codon positions, the strongest effect being observed for the correlation between GC3 content and recombination rates (Spearman’s rank correlation test, all P-values < 2.2×10^−16^; Figure 7). Since this codon position is less impacted by purifying selection, our results are consistent with a joint role of purifying selection and biased gene conversion in shaping nucleotide variation in the green anole genome.

**Figure 7.**
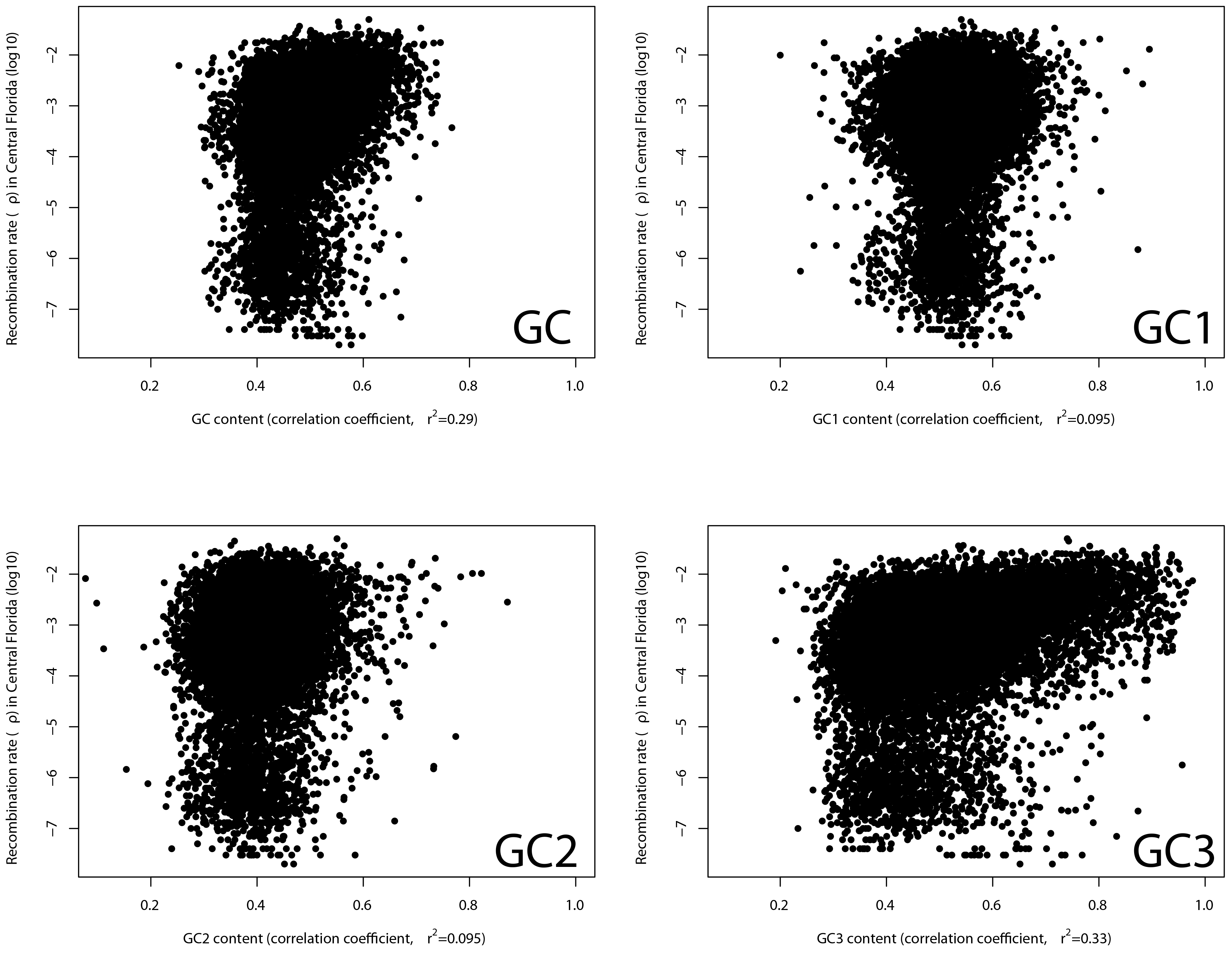
Correlations between p and CDS GC content for all positions, first codon position (GC1), second codon position (GC2) and third codon position (GC3).

## Discussion

### A dynamic demographic history has shaped the genomic landscape of differentiation

Green anole populations are strongly structured and it was hypothesized that successive splits and secondary contact occurred in Florida during the Pleistocene (Tollis and Boissinot 2014; Manthey et al. 2016). Fluctuations in sea level may have generated temporary islands on which isolated populations could have diverged. At last, reconnection of Florida to the mainland would have provided the opportunity for expansion northwards (Soltis et al. 2006). Our results support this claim in three ways. First, splitting times estimated by SMC++ and ∂a∂i suggest a series of splits in Florida between three and two million years ago, a time range during which successions of glacial and interglacial periods may have led to several vicariance events (Lane, 1994; Petuch, 2004). Second, the models receiving the highest support in ∂a∂i were the ones allowing for several events of isolation followed by secondary contact in Florida. Third, we found clear signatures of population expansion in GA and CA at the beginning of the Late Pleistocene, a time when lowering sea levels would have facilitated colonization (Lane, 1994; Petuch, 2004). Despite an old history of divergence, we found clear evidence for gene flow between taxa having diverged in the last two million years. We argue that this makes the green anole a valuable model to study speciation (and its reversal) in the presence of gene flow, as well as identifying genomic incompatibilities and regions under positive selection.

Here, we found evidence of locally reduced diversity due to background selection within Florida in our ∂a∂i models. We note however that models with the highest likelihoods for the Gulf Atlantic-Eastern Florida comparison included heterogeneous migration rates along the genome, and suggested barriers to gene flow limiting introgression from Florida. This could reflect local adaptation through reduced effective migration rates at loci involved in adaptation to northern latitudes (but see (Bierne et al. 2011)).

The role of background selection is further supported by the correlations we observed between diversity, differentiation, and recombination (see below), although we acknowledge that some regions of high divergence and low diversity may have been the targets of positive selection right after population splits (Cruickshank and Hahn 2014). This does not preclude the existence of heterogeneous gene flow along the genome, since we could not properly test the likelihood of models incorporating both of these aspects at once. Instead, this highlights the important role of purifying selection in producing heterogeneous landscapes of differentiation (Cruickshank and Hahn 2014), even in a context of secondary contact where genomic islands resisting gene flow may be more expected.

Recent years have seen a growing interest for the so-called “genomic islands of speciation”, regions that harbor higher differentiation than the genomic background (Feder and Nosil 2010; Ellegren et al. 2012; Nadeau et al. 2012; Wolf and Ellegren 2016). Several studies have since successfully highlighted the important role of heterogeneous migration and selection in shaping diversity in several organisms, such as mussels (Roux et al. 2014), sea bass (Tine et al. 2014) or poplars (Christe et al. 2016). This area of research has however been neglected so far in squamates, preventing any comparison of their genome dynamics at microevolutionary scales with other vertebrates. The green anole is a valuable system to understand local adaptation in reptiles (Campbell-Staton et al. 2017) and the incorporation of our findings in future studies will be valuable to properly test for signals of local adaptation by taking into account the biases induced by demography and the impact of selection at linked sites.

### Unequal diversity between X and autosomal chromosomes suggest a role for selection in accompanying northwards expansion

We detected a significant deviation from a balanced effective sex-ratio in the two populations that recently expanded and colonized North America, with strongly reduced nucleotide diversity on the X chromosome in Gulf Atlantic when compared to autosomal diversity (Sup. Fig. 2). This suggests that the number of females that contributed to the present diversity on the X chromosome may have been extremely reduced compared to the number of males. Since this signature was found only in expanding populations, a possible explanation would be that the colonization of suboptimal habitats (compared to the center of origin in Florida) favored male-biased dispersal. The limited number of available females in the newly colonized regions would have therefore led to a biased sex-ratio in the founding populations and smaller effective population sizes on the X chromosome compared to unbiased expectations.

In *Anolis roquet*, male-biased dispersal is associated with competition, since males disperse more when density increases and competition for females is stronger (Johansson et al. 2008). In *Anolis sagrei*, smaller males tend to disperse more while females are more likely to stay in high quality territories, independently of female density (Calsbeek 2009). The green anole is a polygynous species, with sexual dimorphism and high levels of competition between males (Jenssen et al. 2000). It is therefore likely that competition within sexes may lead to unequal contribution of males and females to the gene pool.

Another non-exclusive possibility lies in recent positive selection on the X chromosome in northern populations. The X chromosome is extremely small compared to autosomes in green anoles, probably not exceeding 20Mb (Rupp et al. 2017). This means that even a few recent selective sweeps would have widespread effects on the entire chromosome, reducing diversity and the effective population size. Since the method implemented in KIMTREE compares estimates of effective population sizes between autosomes and X chromosome, this would result in an artificially biased sex-ratio. Sexual or natural selection may be responsible for this pattern, and our finding calls for further comparisons of sex-biased dispersal and behavior between populations of the green anole. This would give valuable insights on the dynamics of speciation in squamates given the important role of sex chromosomes, for example through the accumulation of Dobzhansky-Muller incompatibilities or divergence at loci involved in mate recognition and choice (Backström et al. 2006; Pryke 2010; Ellegren et al. 2012; Wolf and Ellegren 2016).

### Selection and recombination shape nucleotide composition and diversity at linked sites

We observed strong heterogeneity in recombination rates along the green anole genome. Our results show that this heterogenous recombination landscape plays an important role in shaping genetic diversity in anoles. Both purifying selection and hitchhiking are expected to reduce diversity and increase genetic differentiation (Cruickshank and Hahn 2014). Signatures of selection such as high differentiation and low diversity should be easier to detect in regions of low recombination. Indeed, regions of high diversity that are characterized by high d_XY_ displayed higher recombination rates in the green anole, while regions with high FST were found in regions of low recombination. The lack of clearly marked GC-rich isochores in the green anole genome was imputed to homogeneous recombination rates and possibly weakened or reversed biased gene conversion (Fujita et al. 2011). Our results confirm previous studies (Costantini et al. 2016) claiming that this assumption does not hold upon closer scrutiny.

Biased gene conversion should increase GC content in regions of high recombination (Marais 2003) and this pattern is found across most vertebrate species, including the green anole (Figuet et al. 2014); however, this assumption had never been tested in non-avian reptiles. The strong association that we observed between recombination rate and GC3 content confirms the importance of biased gene conversion for base composition in anoles and more generally squamates.

## Conclusion

*Anolis carolinensis* is an important model organism for biomedical and physiological studies, and benefits from a complete genome sequence that can be used to bridge multiple mechanisms underlying adaptation in natural populations, a key aspect of evolutionary biology (Laland et al. 2011). Genomic resources for anoles are growing and have started uncovering the selective constraints that act on diversity in this clade (Campbell-Staton et al. 2017; Tollis et al. 2018). However, studying the genetic bases of adaptation in anoles cannot be properly addressed without quantifying the patterns that can blur signatures of local adaptation, such as heterogeneous introgression (Roux et al. 2014) or background selection (Hoban et al. 2016). Moreover, the study of intraspecific genetic variation holds promise to address questions at larger evolutionary scale, such as the role of demography, selection and incompatibilities in the process of speciation and genome divergence (Figuet et al. 2014; Romiguier et al. 2014; Seehausen et al. 2014; Chalopin et al. 2015; de la Harpe et al. 2017). Here we sequenced 27 genomes of green anoles and highlighted how secondary contact, expansions, but also heterogeneous recombination and purifying selection have shaped the genomic landscape of differentiation. This study provides a valuable background for precise quantification of the relative importance of selection, demography, and recombination on diversity in non-avian reptiles.

## Methods

### DNA Extraction and Whole Genome Sequencing

Whole genome sequencing libraries were generated from *Anolis carolinensis* liver tissue samples collected between 2009 and 2011 (Tollis et al. 2012), and porcatus and allisoni tissue samples generously provided by Breda Zimkus at Harvard University. For each of the 29 samples, DNA was isolated from ethanol preserved tissue using Ampure bead beads per the manufacturers protocol. Illumina TRU-Seq paired end libraries were generated using 200 ng of DNA per sample and sequenced at the NYUAD Center for Genomics And Systems Biology Sequencing Core (http://nyuad.nyu.edu/en/research/infrastructure-and-support/core-technology-platforms.html) with an Illumina HiSeq 2500. Read quality was assessed with FastQCv0.11.5 (http://www.bioinformatics.babraham.ac.uk/projects/fastqc) and Trimmomatic (Bolger et al., 2014) was subsequently used to remove low quality bases, sequencing adapter contamination and systematic base calling errors. Specifically, the parameters “trimmomatic_adapter.fa:2:30:10 TRAILING:3 LEADING:3 SLIDINGWINDOW:4:15 MINLEN:36” were used. Samples had an average of 1,519,339,234 read pairs, and after quality trimming 93.3% were retained as paired reads and 6.3% were retained as single reads. Sequencing data from this study have been submitted to the Sequencing Read Archive (https://www.ncbi.nlm.nih.gov/sra) under the BioProject designation PRJNA376071.

### Sequence Alignment and SNP Calling

Quality trimmed reads were aligned to the May 2010 assembly of the *A. carolinensis* reference genome (Broad AnoCar2.0/anoCar2; GCA_000090745.1; Alföldi et al., 2011) and processed for SNP detection with the assistance of the NYUAD Bioinformatics Core, using NYUAD variant calling pipeline. Briefly, the quality-trimmed FastQ reads of each sample were aligned to the AnoCar2.0 genome using the BWA-mem short read alignment approach (Li and Durbin, 2009) and resulting SAM files were converted into BAM format, sorted and indexed using SAMtools (Li et al. 2009). Picard was then used to identify insertions, deletions and duplications in the sorted BAM files (http://broadinstitute.github.io/picard/) and evaluated using SAMtools (stats and depth). Alignments contained an average of 204,459,544 reads that passed QC, 97.75% mapping and 91.93% properly paired (Table S1). Each re-sequenced genome was then processed with GATK for indel realignment, SNP and indel discovery and genotyping, following GATK Best Practices (Depristo et al. 2011; Van Der Auwera et al. 2014) (DePristo et al., 2011; Van der Auwera et al., 2013). GATK joint genotyping was conducted for increased sensitivity and confidence, and results were selectively compared to results generated from SAMtools mpileup (Li et al., 2009).Filtering was performed in VCFtools (Danecek et al. 2011), with the following criteria: a 6X minimum depth of coverage per individual, a 15X maximum average depth of coverage, no more than 40% missing data across all 29 samples, a minimum quality score of 20 per site, and a minimum genotype quality score of 20.

### Population structure

To assess genetic structure, we conducted a clustering analysis using discriminant analysis of principal components (DAPC) on a dataset of ~6,500 SNPs with less than 20% missing data and thinned every 10kb to minimize linkage. DAPC first estimates principal components (PC) describing variance in SNP datasets, then performs a discriminant analysis on these PC axes to identify genetic groupings. We retained two principal components and two of the linear discriminants. We also described relationships between individuals with the same dataset using the network algorithm implemented in Splitstree v4 (Huson and Bryant 2006). Lastly, we filtered the SNP dataset to include one million randomly-sampled SNPs present in a minimum of 80% of the individuals for use as input in RAxML v8 (Stamatakis 2014). We used RAxML to create a maximum-likelihood phylogeny, using the GTRGAMMA model of sequence evolution, and 100 rapid bootstraps to assess support for the phylogeny with the highest likelihood.

We further quantified patterns of diversity and the shape of the allele frequency spectrum in each cluster by computing two summary statistics, the average number of pairwise differences (or nucleotide diversity) per bp, and Tajima’s D, for non-overlapping 5kb windows using the software POPGENOME (Pfeifer et al. 2014). We removed windows overlapping ambiguities in the green anole genome using BEDTOOLS v2.25.0 (Quinlan and Hall 2010).

### Demographic estimates without gene flow

We used the multi-epoch model implemented in SMC++ (Terhorst et al. 2016) to reconstruct population size trajectories and time since population split for each of the five genetic clusters of green anoles. This software is an extension of the Pairwise Sequentially Markov Coalescent (Li and Durbin 2011) that uses the spatial arrangement of polymorphisms along genome sequences to naively infer variation in effective population sizes and splitting times between populations. An advantage of this algorithm is that it is phase-insensitive, limiting the propagation of phasing errors that can bias effective population size estimates for recent times (Terhorst et al. 2016). Within each of the 5 genetic clusters, we created one dataset per individual for each of the six autosomes and combined those individual datasets to estimate composite likelihoods. A mutation rate of 2.1×10^−10^ per site per generation and a generation time of one year (Tollis and Boissinot 2014) were assumed to translate coalescence times into years. We also estimated splitting times between Carolinas and Eastern Florida, Gulf Atlantic and Eastern Florida, East Florida and Western Florida, Western and South Florida. Note that splitting times are estimated assuming that no gene flow occurs after the split.

### Effective sex-ratio (ESR)

Sex-biased contribution to the gene pool is a critical aspect of demographic dynamics and is often impacted by variation in social structure between populations. We used the algorithm implemented in KIMTREE (Gautier and Vitalis 2013; Clemente et al. 2018) to estimate branch lengths from our SNP dataset and infer the effective sex-ratios for each of the five genetic clusters. This method is robust to linkage disequilibrium (LD), small sample sizes, and demographic events such as bottlenecks and expansions. To increase the number of usable markers, and since the authors recommend working with recently diverged populations, we focused on the recent northwards colonization, including individuals from the East Florida, Gulf Atlantic, and Carolinas genetic clusters.

Briefly, the method builds a hierarchical Bayesian model to estimate the evolution of SNP frequencies along branches of a population tree provided by the user. Genetic drift along branches is estimated by a time-dependent diffusion approximation. In this framework, branch length τ is proportional to the time since divergence in generations (t) scaled by the effective population size (N_e_), such as τ ≡ t/2N_e_. The method can jointly contrast allele frequencies between autosomal and sex-linked markers to estimate the relative contribution of males and females to each generation (the effective sex-ratio, ESR). The ESR can then be seen as a comparison of the effective population sizes estimates obtained from autosomes and the X chromosome.

We sexed individuals by taking advantage of the expected relationship between depths of coverage at autosomal and sex-linked loci in males and females. Since females are XX and males XY, the latter are expected to display a two-times lower coverage at X-linked sites compared to autosomal loci (Sup. Fig 1). We then adjusted allele frequencies for all X-linked scaffolds, including Linkage Group b (Alföldi et al. 2011) and several scaffolds (GL343282,GL343364,GL343550,GL343423,GL343913,GL343947,GL343338,GL343417) recently identified as belonging to the green anole’s sex chromosome (Rupp et al. 2017). We counted one haplotype per male and two per female. To obtain confidence intervals over ESR estimates, we generated 50 pseudo-replicated datasets by randomly sampling 5,000 autosomal and 5,000 sex-linked SNPs with no missing data. The algorithm was started with 25 pilot runs of 1.0 iterations each to adjust the parameters of the Monte Carlo Markov Chain (MCMC). The MCMC itself was run for 100,000 generations and sampled every 25 iterations after a burn-in of 50.0 iterations. Convergence for all parameters was assessed by visually inspecting posterior sampling in R (R Core team 2016). For each replicate i, we estimated the support for biased sex-ratio (Si) such as:

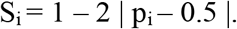

with S_i_ < 0.05 being interpreted as a strong support for biased sex-ratio and where p_i_ is the proportion of posterior MCMC samples with an ESR higher than 0.5.

### Model comparison of realistic demographic scenarios

None of the previous population genetics studies of green anoles have ever precisely quantified the strength nor the timing of gene flow between genetic clusters. We addressed this issue by comparing different demographic scenarios for two pairs of sister clades including at least 11 individuals (EF and GA, EF and WF). We used the diffusion approximation-based likelihood approach implemented in the dadi software (Gutenkunst et al. 2009). We compared a set of scenarios of strict isolation (SI), isolation with migration (IM), ancient migration (AM) with one or two (PAM) periods of gene flow and secondary contact (SC) with one or two (PSC) periods of gene flow (see (Christe et al. 2016) for a detailed summary). We added complexity to this set of basic scenarios by allowing for a combination of population expansion (prefix ‘ex’), heterogeneous asymmetric migration rates (suffix ‘2M2P’) and heterogeneous effective population size (suffix ‘2N’) among loci. These additions were made to incorporate the genome-wide effects of background selection on linked neutral sites (so-called ‘linked selection’) and model genomic islands resisting gene flow (Cruickshank and Hahn 2014). We also tested scenarios with both asymmetric migration rates and heterogeneous population sizes but were unable to reach convergence. Overall, we compared 34 scenarios combining these features, using a set of scripts available on dryad (Christe et al. 2016) and a modified version of ∂a∂i kindly provided by Christelle Fraïsse (available at https://datadryad.org//resource/doi:10.5061/dryad.3bc76 and http://methodspopgen.com/wp-content/uploads/2017/12/dadi-1.7.0modif.zip). We extracted for each pairwise comparison a set of ~12,000 SNPs with no missing data and thinned every 100,000 bp to meet the requirement of independence among loci that is needed to properly compare the composite likelihoods estimated by Sadi. We extracted the unfolded joint sites frequency spectra (SFS) by polarizing alleles using *A. porcatus* and *A. allisoni* as references. We considered ancestral the allele found at a minimal frequency of 75% in those two individuals or found fixed in one of them if the other individual was missing. We note that the a i models include a parameter (O) estimating the proportion of correctly polarized sites. We evaluated each model 30 times and retained the replicate with the highest likelihood for model comparison. Models were compared using the Akaike information criterion (AIC). For the best model, we calculated uncertainties over the estimated parameters using a non-parametric bootstrap procedure, creating 100 pseudo-observed datasets (POD) by resampling with replacement from the SFS. We used the procedure implemented in the dadi.Godambe.GIM_uncert() script to obtain a maximum-likelihood estimate of 95% confidence intervals (Coffman et al. 2016). ∂a∂i parameters are scaled by the ancestral population size N_ref_. For the sake of comparison with SMC++ estimates, parameters were converted into demographic units by estimating the ancestral effective population size as the harmonic mean of the SMC++ estimates before splitting time for all pairs of populations.

### Estimating recombination rates

We used the LDHat software (McVean et al. 2002) to estimate effective recombination rates (ρ=4Nr with r the recombination rate per generation and N the effective population size) along the green anole genome. Unphased genotypes were converted into LDHat format using VCFtools (option–ldhat). Since LDHat assumes that samples are drawn from a panmictic population, we focused on the Eastern Florida clade for which sampling effort was the highest (n=8 diploid individuals). We used precomputed likelihood lookup tables with an effective population mutation rate (0) of 0.001, which was the closest from the 0 value estimated from our dataset (0 ~ 0.004) and used the lkgen module to generate a table fitting the number of observed samples (16 chromosomes). Recombination rates were estimated over 500kb windows with 100kb overlaps using the Bayesian reversible MCMC scheme implemented in the interval module. The chain was run for 1,000,000 iterations and sampled every 5000 iterations with a large block penalty of 20 to avoid overfitting and minimize random noise. The first 100,000 generations were discarded as burn-in. Convergence under these parameters was confirmed by visually inspecting MCMC traces for a subset of windows. We averaged p estimates over nonoverlapping 100kb windows, or over coding sequences (CDS) for subsequent analyses.

### Summary statistics for differentiation and LD

To assess whether selection and low recombination had an effect on diversity and differentiation, we computed two measures of divergence (F_ST_ and d_XY_) over non-overlapping 100kb windows for the three divergent Floridian lineages. Comparison between those two statistics for a given genomic region has been proposed as a way to disentangle the effects of gene flow and selection (Cruickshank and Hahn 2014). As a sanity check, we computed the ZZ statistics (Rozas et al. 2001) to assess whether LDHat estimates of ρ were consistent with the genomic distribution of LD. This statistic contrasts LD between adjacent pairs of SNPs to LD calculated over all pairwise comparisons in a given window. High values are suggestive of increased intragenic recombination. All statistics were computed in the R package POPGENOME (Pfeifer et al. 2014).

### GC content

We extracted CDS sequences for all green anole genes from the ENSEMBL database (available at ftp://ftp.ensembl.org/pub/release-88/fasta/anoliscarolinensis/cds/). For each CDS, we estimated overall GC content, as well as GC content at first, second, and third codon position (GC1, GC2, GC3) using the R package seqinr (Charif et al., 2015). We used BEDTOOLS (Quinlan and Hall 2010) to extract ρ estimates overlapping exons for each CDS, and averaged ρ over the total CDS length. Spearman’s rho coefficients for correlations between GC content and recombination rates were estimated in R.

## Acknowledgements

We are grateful to Breda Zimkus from the Museum of Comparative Zoology Cryogenic Collection in Harvard and J. Rosado from the Herpetology Collection for providing the samples. We also thank Christelle Fraïsse for providing tutorials and the modified version of ∂a∂i that was needed to compare demographic models. We thank Justin Wilcox for his comments on the manuscript. We thank Marc Arnoux from the Genome Core Facility at NYUAD for assistance with genome sequencing. This research was carried out on the High Performance Computing resources at New York University Abu Dhabi. This work was supported by New York University Abu Dhabi (NYUAD) research funds AD180 (to S.B.). The NYUAD Sequencing Core is supported by NYUAD Research Institute grant G1205-1205A to the NYUAD Center for Genomics and Systems Biology.

## Supplementary Table

Table S1. Samples origin, sequencing depth and quality statistics.

### Supplementary Figures

Figure S1: Plot of average depth of coverage for sex-linked markers v. autosomal markers in all 27 green anoles used in this study. Males should fall on the line y=2*x due to the representation bias expected in XY individuals. Females are XX and should fall on the line y=x.

Figure S2: Boxplots of nucleotide diversity across non-overlapping 5kb windows at autosomes and sex-linked scaffolds for three EF females and three Gulf Atlantic females. The analysis was restricted to females to account for haplodiploidy at sex-linked scaffolds. The dotted lines delimit autosomes from sex-linked scaffolds.

